# Profiling metabotropic glutamate receptor 7 expression in Rett syndrome: consequences for pharmacotherapy

**DOI:** 10.1101/2025.05.08.652716

**Authors:** Sheryl Anne D. Vermudez, Geanne A. Freitas, Mackenzie Smith, Rocco Gogliotti, Colleen M. Niswender

## Abstract

We have reported that levels of metabotropic glutamate receptor 7 (mGlu_7_) are dramatically decreased in brain samples from Rett syndrome patients carrying truncation mutations in the *Methyl-CpG Binding Protein 2* (*MECP2*) gene. Additionally, we identified decreases in mGlu_7_ levels in *Mecp2^+/-^* female mice and demonstrated that administration of a positive allosteric modulator (PAM) with activity at mGlu_7_ corrected deficits in cognitive, social, and respiratory domains. Here, we expanded our studies to a larger cohort of RTT samples covering a range of mutations and evaluated expression of the three widely expressed group III mGlu receptors (mGlu_4,7 and 8_). We found significant decreases in mGlu_7_, but not mGlu_4_ or mGlu_8_, mRNA expression across this larger cohort; additionally, we identified a previously unknown and robust correlation in the expression of mGlu_4_ and mGlu_8_ in control individuals. Stratification of RTT patients into individuals with mutations that are clinically correlated with severe versus mild disease revealed statistically significant decreases in mGlu_7_ expression only in patients with mutations that induce more severe symptoms. We then administered the PAM VU0422288 to mice modeling the mild R306C mutation (*Mecp2^R306C/+^*) and found a significant reduction in apneas induced by VU0422888 administration despite no decreases in mGlu_7_ expression in the brainstem or cortex. These results provide the first evidence of potentially broad utility for mGlu_7_ PAMs in reducing apneas in RTT patients.

## Introduction

Rett syndrome (RTT) is caused by mutations in the X-linked Methyl-CpG Binding Protein 2 (*MECP2*) gene, a transcriptional regulator of both local and global gene expression patterns (Amir et al., 1999; Chahrour et al., 2008; Ben-Shachar et al., 2009). RTT patients develop normally during their first 6 to 18 months of life and then undergo a rapid developmental regression, resulting in impairments in language and motor skills and the emergence of breathing abnormalities (i.e., apneas) and seizures (Neul et al., 2010; Gold et al., 2024; Lopes et al., 2024). Given the monogenic nature of the disease, many therapeutic strategies are aimed specifically at MeCP2 and include gene therapy, DNA and RNA editing, X-chromosome reactivation, and pharmacological read-through approaches (reviewed in (Percy 2016; Palmieri et al., 2023; Gold et al., 2024; Lopes et al., 2024); NCT05898620, NCT06152237). However, many of these efforts are complicated by the presence of a narrow therapeutic window, whereby even modest overexpression MeCP2 also impairs neurodevelopment and cell function (Collins et al., 2004; Ramocki et al., 2010; Collins and Neul 2022). One therapeutic strategy that may avoid these confounds is to develop or repurpose drugs that modulate downstream targets of MeCP2. Currently, the sole FDA approved symptomatic treatment for RTT is the compound trofinetide, which is a modified version of a glycine-proline-glutamate tripeptide cleaved from insulin growth factor 1 (IGF-1, (Furqan 2023; Harris 2023; Neul et al., 2023)).

The concerted effort to biobank brain samples from RTT patient autopsies and render broad availability of expression profiling data widely available provides an opportunity to identify disruptions in downstream targets, allowing discovery efforts to begin from a place of translational relevance. We previously performed RNA and protein expression profiling studies from the motor cortex and cerebellum of a cohort of autopsy samples from patients diagnosed with RTT and corresponding age, postmortem interval, and sex-matched controls (Gogliotti et al., 2016; Gogliotti et al., 2017; Gogliotti et al., 2018). These studies identified a number of G protein-coupled receptors (GPCRs) that were changed in expression in RTT patients, including several metabotropic glutamate (mGlu) and muscarinic acetylcholine receptors. One target identified was the mGlu_7_ receptor, which was reduced by approximately 70% in both the cortex and cerebellum of RTT patients with the MeCP2 truncation mutations R168X, R255X, and R270X (Gogliotti et al., 2017). We then showed that abnormal social, respiratory (e.g., apneas), and cognitive phenotypes could be corrected by administering a positive allosteric modulator (PAM) with mGlu_7_ activity, termed VU0422288, to *Mecp2^-/y^* male and *Mecp2^+/-^* female mice (Jalan-Sakrikar et al., 2014; Gogliotti et al., 2017). mGlu_7_ belongs to the group III mGlu receptors, which also includes the related and widely expressed mGlu_4_ and mGlu_8_ receptors (Niswender and Conn 2010), and VU0422288 potentiates the activity of all of these receptor isoforms (Jalan-Sakrikar et al., 2014). In our studies validating mGlu_7_ as a potential target in RTT, we also employed the PAM ADX88178 (Le Poul et al., 2012; Kalinichev et al., 2014), which potentiates the activity of mGlu_4_ and mGlu_8_ but not mGlu_7_; this compound was ineffective in correcting phenotypes (Gogliotti et al., 2017). As an additional validation that mGlu_7_ was the relevant target, we also showed blockade of the beneficial effects of VU0422288 with a highly selective mGlu_7_ negative allosteric modulator (NAM), ADX71743 (Kalinichev et al., 2013; Gogliotti et al., 2017).

Our findings in RTT correlate with emerging reports that primary loss-of-function mutations within the *GRM7* gene cause seizures, developmental delay, autism spectrum phenotypes, ADHD, autism, hypomyelination, and intellectual impairments (Sanders et al., 2012; Liu et al., 2015; Charng et al., 2016; Reuter et al., 2017; Fisher et al., 2018; Marafi et al., 2020; Fisher et al., 2021; Jdila et al., 2021; Song et al., 2021; Freitas and Niswender 2023), as well as reports that polymorphisms in *GRM7* are linked to diseases such as epilepsy, cognitive impairments, and ADHD (Mick et al., 2008; Ohtsuki et al., 2008; Ganda et al., 2009; Shibata et al., 2009; Saus et al., 2010; Alliey-Rodriguez et al., 2011; Breen et al., 2011; Hamilton 2011; Elia et al., 2012; Yang and Pan 2013; Kandaswamy et al., 2014; Park et al., 2014; Haenisch et al., 2015; Jajodia et al., 2015; Nho et al., 2015; Noroozi et al., 2019). One of these clinical *GRM7* mutations affects the trafficking and stability of the receptor and several of these mutations result in severe impairments in neuronal maturation and myelination (Fisher et al., 2021; Song et al., 2021). Overall, these data suggest that low levels of mGlu_7_ are deleterious and establish the hypothesis that mGlu_7_ activation or potentiation using small molecule PAMs could represent a novel therapeutic strategy for various phenotypes seen in neurodevelopmental syndromes, including RTT.

Our original studies with mGlu_7_ profiling examined expression in patients with truncation mutations in MeCP2; however, many RTT patients have missense mutations in the gene that result in different degrees of disease severity (Weaving et al., 2003; Bebbington et al., 2008; Neul et al., 2008; Cuddapah et al., 2014; Townend et al., 2018; Collins and Neul 2022; Downs et al., 2024). The availability of additional autopsy brain samples from patients with RTT syndrome at the University of Maryland and Harvard Brain Banks has offered us an opportunity to profile additional patients, including those with missense mutations, rather than truncation mutations in *MECP2,* for mGlu_7_ expression. The latter point is salient as the type and location of each mutation is predictive of clinical severity (Yamashita et al., 2001; Leonard et al., 2003; Bebbington et al., 2008; Renieri et al., 2009; Cuddapah et al., 2014; Collins and Neul 2022). For example, the R133C and R306C mutations, along with late truncating mutations, present with milder symptoms, while early truncating mutations, such as R168X and R255X, result in more severe phenotypes (Yamashita et al., 2001; Leonard et al., 2003; Bebbington et al., 2008; Renieri et al., 2009; Cuddapah et al., 2014). Since our original characterization was conducted using samples with early truncating mutations, it is unknown whether mGlu_7_ is disrupted in patients with mild missense mutations. The potential for mutation-specific expression patterns is important, as it raises the possibility that mGlu_7_ PAMs could be most effective in patients with more severe mutations and less effective in other populations and could inform inclusion criteria for potential clinical trials. Here, we profiled the expression of mGlu_7_, along with the related mGlu_4_ and mGlu_8_ receptors, in brain samples derived from the temporal cortex from a 14-sample cohort of controls and a 41-sample cohort of RTT patients genotyped for *MECP2* mutations. We also profiled the expression of the other widely expression group III mGlu receptors, mGlu_4_ and mGlu_8_ (Gogliotti et al., 2017). Across the group, mGlu_7_ mRNA levels were significantly reduced between control and RTT individuals but mGlu_4_ and mGlu_8_ levels were not. When these data were separated by mutations that correlate with mild versus severe forms of RTT, we found significantly decreased mGlu_7_ expression in cortical samples from RTT patients in individuals with mutations that result in more severe forms of RTT.

The observation that some RTT patients do not exhibit low levels of mGlu_7_ suggests that an mGlu_7_ PAM may not be as effective in these individuals. Here, we used a mouse model of RTT of the milder clinical mutation, R306C, and find that mGlu_7_ expression levels are not reduced in most brain regions, including the brainstem, from these animals; the exception was the hippocampus, where we did observe a significant reduction. We then treated *Mecp2^+/+^* and *Mecp2^R306C/+^* female mice with a 30 mg/kg dose of the PAM VU0422288, which was previously shown to be effective in *Mecp2^+/-^* mice (Gogliotti et al., 2017), and found that VU0422288 still exhibited efficacy in reducing apneas in *Mecp2^R306C/+^* animals without affecting other breathing parameters. These results provide the first evidence validating the efficacy of mGlu_7_ PAMs in disease models of RTT that result from mutations that induce milder phenotypes.

## Materials and Methods

### Human Brain Sample Profiling

Human samples were obtained from the National Institutes of Health NeuroBioBank (neurobank.nih.gov) under Public Health Service contract HHSN-271-2013-00030. The tissues were post-mortem and fully de-identified, and as such are classified as exempt from human subject research regulations. The Brodmann Area (BA) 38 (temporal cortex) samples used for this study are shown in Table 1. For the majority of these samples, the *MECP2* mutation was unknown upon receipt, and we performed TOPO TA cloning and Sanger sequencing to identify the *MECP2* mutation for each sample using primers that span the length of and corresponding to the e1 isoform (predominantly expressed in the brain) of the gene.

**Table 1.** Demographics and group III mGlu receptor mRNA expression for human patient samples. mRNA assessments were performed in duplicate and normalized to *G6PD* expression.

| Controls | UNMB number | Age | PMI | *MECP2*<br>Mutation | *GRM7*<br>mRNA | *GRM4*<br>mRNA | *GRM8*<br>mRNA |
| --- | --- | --- | --- | --- | --- | --- | --- |
| CTL-1 | 1038 | 24 | 7 | N/A | 65 | 128 | 120 |
| CTL-3 | 5161 | 10 | 22 | N/A | 158 | 113 | 113 |
| CTL-4 | 5554 | 13 | 15 | N/A | 119 | 105 | 109 |
| CTL-6 | 5844 | 42 | 12 | N/A | 23 | 56 | 65 |
| CTL-7 | 5566 | 15 | 23 | N/A | 115 | 114 | 122 |
| CTL-8 | 5309 | 14 | 8 | N/A | 120 | 120 | 111 |
| CTL-9 | 4330 | 19 | 19 | N/A | 209 | 82 | 77 |
| CTL-10 | 5446 | 18 | 18 | N/A | 114 | 121 | 118 |
| CTL-11 | 5538 | 19 | 24 | N/A | 104 | 99 | 93 |
| CTL-12 | 6544 | 30 | 23 | N/A | 124 | 98 | 101 |
| CTL-13 | 5646 | 21 | 23 | N/A | 100 | 87 | 92 |
| CTL-14 | 5669 | 25 | 29 | N/A | 77 | 137 |  |
| CTL-15 | 5670 | 17 | 22 | N/A | 39 | 34 |  |
| CTL-16 | 5751 | 25 | 24 | N/A | 34 | 111 | 82 |
| **AVE** |  | **20.9** | **19.2** |  | **100** | **100** | **100** |
| **SD** |  | **8.2** | **6.5** |  | **50.2** | **28.0** | **18.5** |
| **SEM** |  | **2.2** | **1.7** |  | **13.4** | **7.5** | **5.3** |
| RTT | Maryland BB<br>number | Age | PMI | *MECP2*<br>Mutation | *GRM7*<br>mRNA | *GRM4*<br>mRNA | *GRM8*<br>mRNA |
| Rett-1 | AN19242 | 41 |  | R255X | 75 | 151 | 158 |
| Rett-3 | AN15579 | 11 | 9.6 | R255X | 43 | 148 | 158 |
| Rett-4 | AN04573 | 12 | 5.0 | R270X | 165 | 109 | 109 |
| Rett-5 | AN08016 | 8 | 2.9 | R255X | 86 | 109 | 155 |
| Rett-6 | AN02091 | 24 | 15.8 | R255X | 36 | 98 | 131 |
| Rett-7 | AN04121 | 10 | 23.5 | R270X | 83 | 137 | 60 |
| Rett-8 | AN01193 | 23 | 10.5 | Mut neg | 1481* | 133 | 88 |
| Rett-11 | AN01508 | 26 | 28.1 | R306C | 96 | 618* | 76 |
| Rett-12 | AN18782 | 28 | 33.4 | Unknown | 45 | 97 | 82 |
| Rett-13 | AN13266 | 33 | 23.7 | R270X | 76 | 83 | 89 |
| Rett-14 | AN17730 | 22 | 31.8 | P302L | 24 | 69 | 106 |
| Rett-15 | AN05282 | 22 | 18.1 | Unknown | 128 | 112 | 111 |
| Rett-16 | AN15264 | 35 | 25.4 | R168X | 52 | 78 | 104 |
| Rett-17 | AN00811 | 31 | 30.0 | Mut neg | 32 | 132 | 59 |
| Rett-18 | AN15541 | 28 | 12.1 | Unknown | 102 | 71 | 80 |
| Rett-19 | AN07799 | 25 | 31.0 | T158M | 63 | 170 | 141 |
| Rett-20 | AN13274 | 26 | 38.5 | Mut neg | 45 | 149 | 90 |
| Rett-21 | AN06847 | 24 | 24.6 | T158M | 51 | 97 | 79 |
| Rett-22 | AN05652 | 41 | 48.8 | R133C | 162 | 78 | 98 |
| Rett-23 | AN07309 | 12 | 16.8 | R255X | 204* | 140 | 50 |
| Rett-24 | AN05180 | 57 | 16.7 | CTD >398 | 112 | 83 | 137 |
| Rett-25 | AN01082 | 7 | 33.2 | Y141X | 118 | 74 | 116 |
| Rett-26 | AN17849 | 12 | 3.1 | R168X | 73 | 137 | 137 |
| Rett-27 | AN14876 | 26 | 17.1 | R270X | 28 | 58 | 115 |
| Rett-28 | AN05375 | 20 | 23.3 | G269del | 51 | 88 | 64 |
| Rett-29 | AN07094 | 18 | 4.4 | Mut neg | 29 | 124 | 36 |
| Rett-30 | AN13427 | 33 | 17.0 | L386fs | 11 | 21 | 26 |
| Rett-31 | AN15255 | 62 | 24.0 | Mut neg | 42 | 78 | 80 |
| Rett-32 | AN05266 | 21 | 19.7 | R168X | 59 | 66 | 66 |
| Rett-33 | AN09497 | 27 | 32.1 | R255X | 73 | 162 | 190 |
| Rett-34 | AN19272 | 16 | 27.6 | A2V | 22 | 27 | 11 |
| Rett-35 | AN02949 | 15 | 21.3 | T158M | 47 | 180 | 212 |
| Rett-36 | AN01220 | 10 | 24.0 | M246del | 479* | 121 | 86 |
| Rett-37 | AN05357 | 12 | 8.1 | R168X | 26 | 112 | 201 |
| Rett-38 | AN03862 | 35 | 32.5 | G269del | 51 | 111 | 96 |
| Rett-39 | AN06744 | 29 | 17.0 | Mut neg | 24 | 110 | 110 |
| Rett-41 | AN12551 | 20 | 14.1 | Mut neg | 55 | 398* | 107 |
| Rett-43 | AN08774 | 6 | 23.5 | P322L | 73 | 153 | 167 |
| Rett-45 | AN07559 | 8 | 22.7 | R306C | 85 | 179 | 34 |
| Rett-46 | AN01325 | 8 | 12 | Mut neg | 36 | 126 | 131 |
| Rett-48 | 8870 |  |  | Mut neg | 52 | 220 | 160 |
| **AVE** |  | **23.1** | **21.0** |  | **64.0** | **112.6** | **105.0** |
| **SD** |  | **12.9** | **10.2** |  | **37.0** | **42.2** | **46.8** |
| **SEM** |  | **2.0** | **1.6** |  | **13.4** | **6.8** | **7.3** |

PMI: postmortem interval. mGlu<sub>7</sub> dimer and monomer bands were quantified for each sample. \*These samples are outliers by ROUT analysis.

### Animals

All animals used in the present study were group housed with food and water given ad libitum and maintained on a 12hr light/dark cycle. Animals were cared for in accordance with the National Institutes of Health Guide for the Care and Use of Laboratory Animals. All studies were approved by the Loyola Institutional Animal Care and Use Committee and took place during the light phase. *Mecp2^306C/+^* (B6.129P2(C)-*Mecp2^tm5.1Bird^*/J, stock no. 026847) were obtained from The Jackson Laboratory (Bar Harbor, ME, USA) and maintained on a C57BL/6J background by breeding *Mecp2^306C/+^* with WT C57BL/6J mice (The Jackson Laboratory, stock no. 000664). As a reflection of the predominantly female RTT patient population, female *Mecp2^306/+^* mice were utilized and aged to at least 20 weeks of age prior to experiments.

### Total RNA preparation and quantitative RT-PCR

As previously reported (Smith et al., 2022), approximately 1 g of the temporal cortex was impact-dissociated under dry ice and then pulverized using mortar and pestle under liquid nitrogen. Total RNA was prepared from 200 mg of tissue using standard trizol-chloroform methodology. cDNA from 2μg of total RNA was synthesized using a SuperScript™ VILO™ cDNA Synthesis Kit (ThermoFisher, cat. no. 11754050). qRT-PCR (CFX96, Bio-Rad, Vanderbilt University Medical Center MCBR Core) on 50ng/9μL cDNA was run in duplicate using TaqMan™ Fast Universal PCR Master Mix (2X), no AmpErase™ UNG (Life Technologies, cat. no. 4352042) and Life Technologies gene expression assays for human *GRM4* (Hs00904641_m1), *GRM7* (Hs00990476), *GRM8* (Hs00945353_m1) and glucose 6-phosphate dehydrogenase (*G6PD,* Hs00166169_m1). Ct values for each sample were normalized to *G6PD* expression and analyzed using the delta–delta Ct method as performed in (Gogliotti et al., 2017). Values exceeding two times the standard deviation were classified as outliers. Each value was compared to the average delta-Ct value acquired for control human samples and calculated as percent-relative to the average control delta-Ct.

### Total and Synaptosomal Protein Preparation and Western Blotting

The cortex, hippocampus, striatum, cerebellum and brainstem were microdissected from 28-week-old female WT (*Mecp2^+/+^*) and *Mecp2^306C/+^* littermates. Synaptosomes were isolated according to a protocol described in (Fentress et al., 2013). Mouse (50μg) total protein was electrophoretically separated using a 4-20% SDS polyacrylamide gel and transferred onto a nitrocellulose membrane (iBlot2, ThermoFisher). Membranes were blocked in TBS Odyssey blocking buffer (LI-COR) for 1hr at room temperature. Membranes were probed with primary antibodies overnight at 4°C: rabbit anti-mGlu_7_ (1:1000, Millipore/Upstate cat no. 07-239), mouse anti-tubulin (1:5000, Abcam cat. no. ab44928) or mouse anti-Gapdh (1:5000, ThermoFisher, cat. no. MA5–15738), followed by the fluorescent secondary antibodies: goat anti-rabbit (800nm, 1:5000, LI-COR, cat. no. 926– 32211) and goat anti-mouse (680nm, 1:10,000, LI-COR, cat. no. 926–68020). Fluorescence was detected using the Odyssey (LI-COR) imaging system at the Vanderbilt University Medical Center Molecular Cell Biology Resource (MCBR) Core and then quantified using the Image Studio Lite software (LI-COR). Levels of monomeric and dimeric mGlu_7_ were normalized to tubulin.

### Drugs

VU0422288 was synthesized at the Warren Center for Neuroscience Drug Discovery according to published methods (Jalan-Sakrikar et al., 2014).

### Whole Body Plethysmography

Unrestrained 28-week-old *Mecp2^+/+^* and *Mecp2^R306C/+^* mice were placed in a whole-body plethysmograph recording chamber (Buxco, 2-site system) with a continuous inflow of air (1 L/min). Following a habituation period of 30 min, a baseline recording was established for 30 min. Mice were then removed from the chamber, injected with VU0422288 or vehicle, and reacclimated for 30 min, and respiratory measurements were made for an additional 30 min. Analysis was performed using FinePointe Research Suite (v2.3.1.9). Apneas, defined as pauses spanning 2 × the average expiratory time of the previous 2 min, were quantified using the FinePointe apnea software patch, followed by manual spot-checking of the larger data set. Only periods of motion-free recording were analyzed. All filters were applied while the researchers were blinded to the genotype and treatment group.

### Statistical Analysis

Statistics were carried out using GraphPad Prism and Excel (Microsoft). All data shown represent mean ± SEM, sample size is denoted as “n” and the statistical test used for each set of data is described in the respective legend. Statistical significance between genotypes was determined using Student’s t-test or 1-way ANOVA with post-hoc test indicated in the figure legend.

## Results

We previously profiled mGlu_7_ expression using motor cortical samples from seven RTT patients with either R168X, R255X, or R270X mutations in MeCP2 as well as eight age, brain region, post-mortem interval, and sex matched controls (Gogliotti et al., 2017). These experiments identified a significant reduction in expression of the mGlu_7_ receptor in the cortex of RTT patients. Here, we expanded these studies to samples from the temporal cortex of a larger, 41 sample, cohort of RTT patients representing a range of *MECP2* mutations (Table 1 lists each patient’s mutation, post-mortem interval, and age). These mutations ranged from the methyl binding domain (Y141X, T158M, R168X), to the transcription repression domain (R255X, G269Del, R270X), to the Nuclear hormone receptor Co-Repressors (NCoR) interaction domain/C-terminal domain (P302L, R306C, P322L, CTD>398). Within our cohort, there were six samples for which we did not detect a mutation in *MECP2* by Sanger sequencing, but the patient had received a clinical RTT diagnoses. Three additional samples were ranked as “unknown” due to poor sequencing results; these patients had also all received a RTT diagnosis.

There was no difference between control and RTT subjects for either age (20.9 ± 2.2 years, control, versus 23.1 ± 2.0 years, RTT, p=0.5404, unpaired t-test, Mean ± SEM,) or postmortem interval (19.2 ± 1.7, control versus 21.0 ± 1.6 hours, RTT, p=0.4684, unpaired t-test, Mean ± SEM) between the control and RTT cohorts.

We performed qRT-PCR to examine *GRM7* mRNA expression (Table 1, Figure 1) across this cohort of control and RTT patients. Expression values in three RTT samples were ranked as outliers in this dataset and these samples were removed from the analysis in Figure 1 but are noted in Table 1. We observed a statistically significant (**p=0.0067, unpaired t-test between control and RTT) reduction in *GRM7* mRNA expression across the cohort (Figure 1A, black versus red, Table 1). We performed similar profiling for mRNA encoding *GRM4* and *GRM8*, the other widely CNS-expressed group III mGlu receptors, and we did not detect significant differences between control and RTT patients for mRNAs encoding these receptors (Figure 1B, C, Table 1; *GRM4*, p=0.3047, *GRM8*, p=0.7170, unpaired t-test).

**Figure 1.**
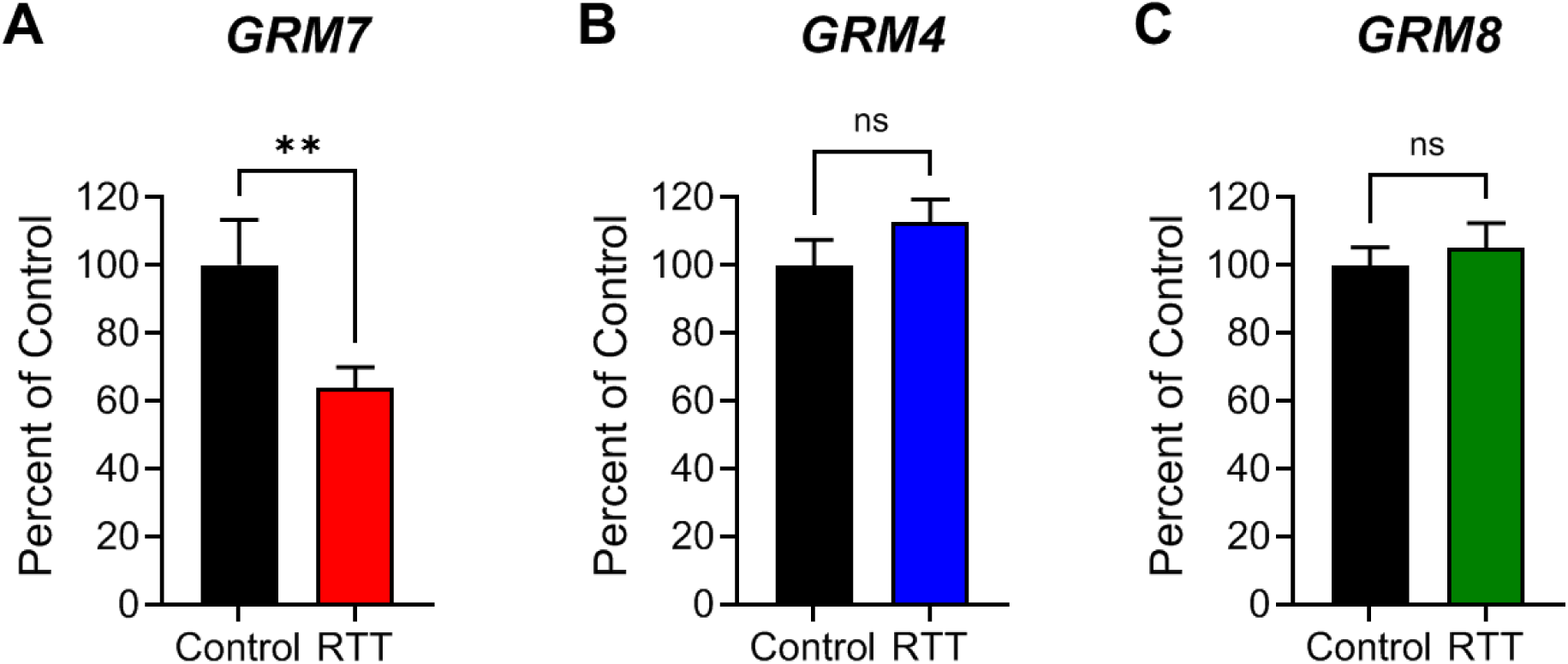
Among the group III mGlu receptors, only *GRM7* mRNA is decreased in cortical autopsy samples from RTT patients. RNA was prepared from human tissue from the BA38 region of the temporal cortex and quantitative RT-PCR was performed for (A) *GRM7*, (B) *GRM4*, and (C) *GRM8*. Data shown are mean ± SEM and were performed in duplicate and averaged. Data were normalized to *G6PD* control and analyzed as unpaired t-test for each receptor (*GRM7*: control versus RTT, p=0.0012 (**); *GRM4* and *GRM8*: ns). Outliers in each dataset are noted in Table 1.

To determine if there was a correlation of group III mGlu mRNA expression with mutation location or disease severity, we next binned samples into mild and severe groups and also included samples that were ranked as *MECP2* mutation-negative. The mild group contained samples corresponding to the mutations R133C, T158M, P302L, R306C, and CTD>398, and the severe group included samples corresponding to the mutations R168X, R255X and R270X; we confirmed that there were no significant differences between control and this subset of samples in terms of age (20.9 ± 2.2 control versus 24.3 ± 2.7 years, RTT, p=0.4116, unpaired t-test) and PMI (19.2 ± 1.7 control versus 21.9 ± 1.9 hours, RTT, p=0.3780, unpaired t-test). These studies revealed statistically significant reductions in *GRM7* mRNA in patients with mutations in the severe but not the mild group (Figure 2A, Table 1). Interestingly, the group of patients for which no *MECP2* mutation was identified showed a very tight cluster of significant reduced *GRM7* mRNA expression (Figure 2A, Table 1). Separating the samples into mild, severe, and mutation negative groups did not result in any statistically significant differences in expression for *GRM4* or *GRM8* (Figure 2B and C). We then performed correlation analyses to determine if there was co-regulation of *GRM7* with either *GRM4* or *GRM8* in the human cortex (Figure 3), as the group III mGlu receptors have been shown to heterodimerize (Doumazane et al., 2011; Liu et al., 2017; Moreno Delgado et al., 2017; Xiang et al., 2021; Lin et al., 2022). These studies revealed that there was no correlation between *GRM7* with *GRM4* or *GRM8* expression (Figure 3A, B). Interestingly, however, there was a highly significant correlation in expression between *GRM4* and *GRM8*, particularly in the control samples (***p=0.0004; Figure 3C) and also in the RTT cohort (**p=0.0027, Figure 3D). To our knowledge, this expression correlation has not been reported for these two receptors in the human brain. Finally, we performed correlations of *GRM7*, *GRM4*, and *GRM8* with post-mortem interval (PMI) and age of the samples in our cohort. No correlation was established for any of the three transcripts with PMI (data not shown); however, we quantified a negative correlation of *GRM7* and *GRM8* with age in control, but not RTT samples (*GRM7*, R^2^ = 0.30, *p = 0.04; *GRM8*; R^2^ = 0.42, *p = 0.02).

**Figure 2.**
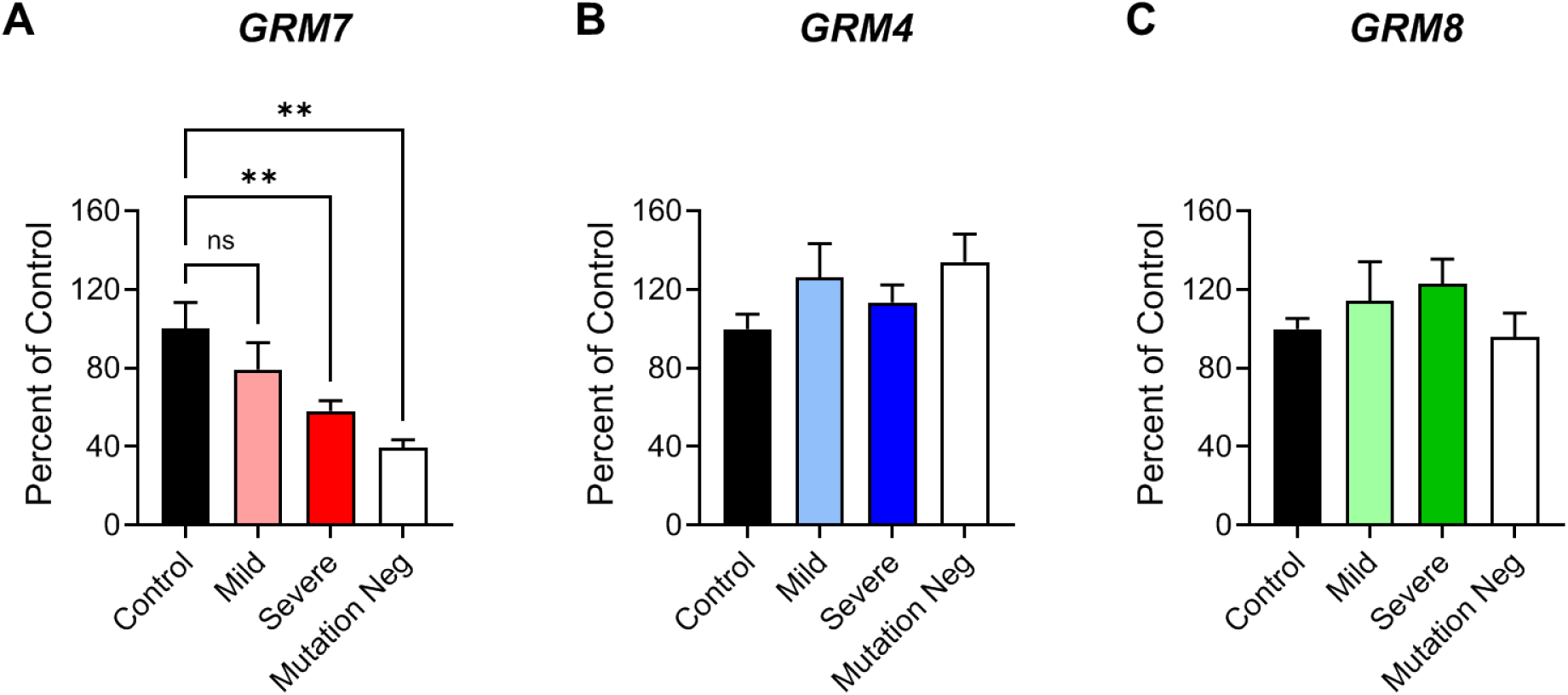
*GRM7* expression is significantly decreased in patients with mutations that induce more severe, but not mild, RTT symptoms. (A) *GRM7* mRNA is significantly reduced in patients with severe (R168X, R255X, R270X) mutations (red) and in patients with mutation-negative RTT (green), but not in patients with mild mutations (R133C, T158M, and R306C). (B) *GRM4* and (C) *GRM8* are not significantly different in any group versus control. Data represent mean +/- SEM and were analyzed using one-way ANOVA; for *GRM7*, F (3,41) 5.928, **p=0.0019; Dunnett’s post-hoc test, control versus mild, p=0.4064; control versus severe, **p=0.0095, control versus mutation negative, **p=0.0013. *GRM4* one-way ANOVA, F (3, 40) 1.783, p=0.1659. *GRM8* one-way ANOVA, F (3, 39) = 1.098, p=0.3617.

**Figure 3.**
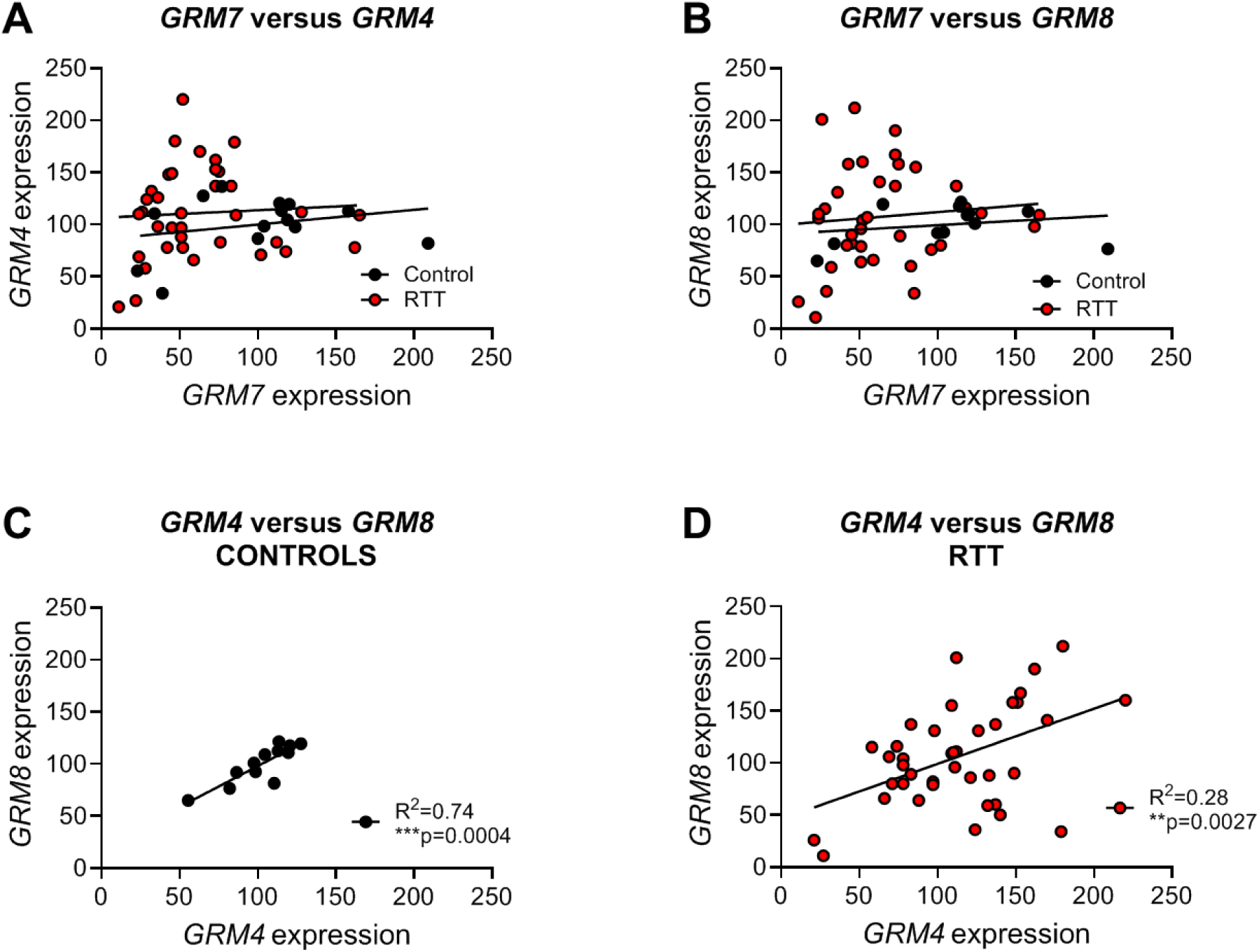
*GRM7* mRNA expression does not correlate with either *GRM4* and *GRM8*; however, *GRM4* and *GRM8* expression exhibits a significant correlation. mRNA expression data from the experiments in Table 1, Figure 1, and Figure 2 were correlated by linear regression for (A) *GRM7* and *GRM4*, (B) *GRM7* and *GRM8*, and *GRM4* and *GRM8* for control (C) and (D) RTT samples.

The observation that reductions in mGlu_7_ were not present in every patient suggests that RTT patients with certain mutations may not respond to a PAM with mGlu_7_ activity. To address this possibility, we chose the R306C mouse model of RTT and profiled synaptic mGlu_7_ protein expression in multiple brain areas from 28-week-old female mice. Expression studies showed a significant decrease in mGlu_7_ protein levels in the hippocampus but not in other brain areas such as the cortex, cerebellum, or brainstem (Figure 4, Supplementary Figures 1-4). Our previous work has shown that VU0422288 robustly reduces apneas in *Mecp2^+/-^* animals as assessed using whole body plethysmography (Gogliotti et al., 2017). In *Mecp2^306C/+^* animals, we also observed a significant apnea reversal when the animals were administered a 30 mg/kg dose of VU0422288 (Figure 5A), but no change in breath rate or minute volume (Figures 5B and 5C). This result was observed despite the normal levels of mGlu_7_ in the brainstem, suggesting that VU0422288’s efficacy may not be linked solely to target disruption, and suggests that mGlu_7_ PAMs have the potential to retain utility across multiple RTT patient subpopulations.

**Figure 4.**
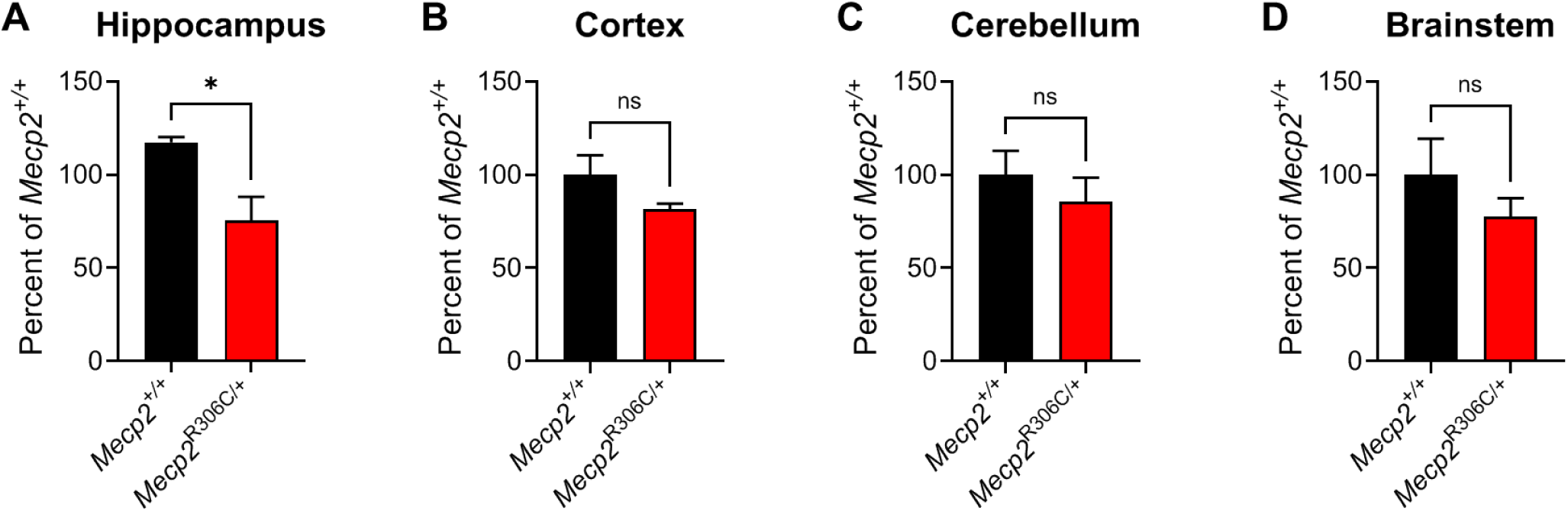
In mice modeling the milder R306C mutation, mGlu_7_ expression in synaptosomes is significantly reduced in the hippocampus, but not the cortex, cerebellum, or brainstem. mGlu_7_ protein levels were assessed in (A) hippocampus, (B) cortex, (C) cerebellum, and (D) brainstem from *Mecp2^+/+^* (WT, black) and *Mecp2^R306C/+^* (red) female mice. Mean +/- SEM shown; *p=0.0249 by unpaired Student’s t-test. Full Western blots are shown in Supplementary Figures 1-4. N=5 per genotype.

**Figure 5.**
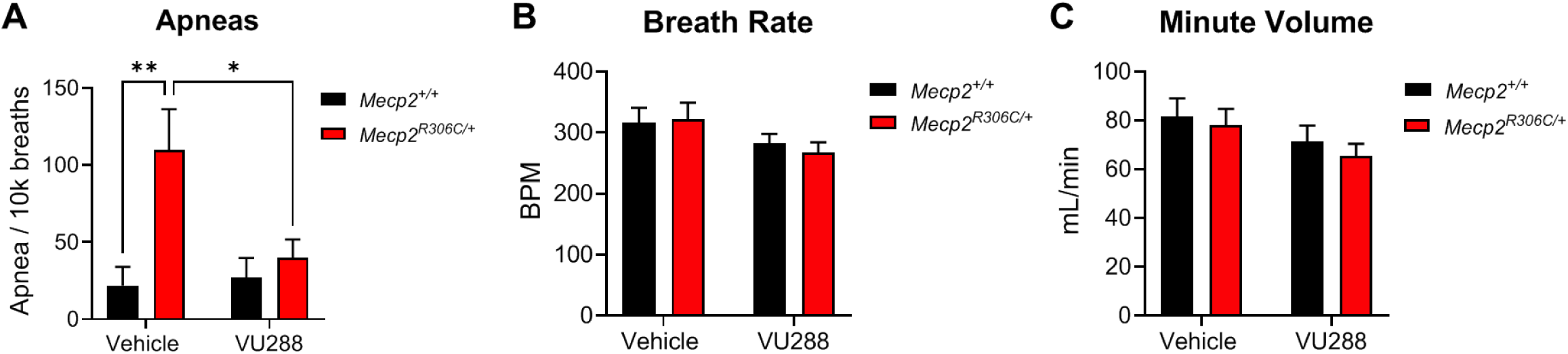
VU0422288 reverses apneas in *Mecp2^R306C/+^* mice. (A) Apneas were assessed using whole body plethysmography (WBP) and normalized to 10,000 breaths for *Mecp2^+/+^* (WT, black) and *Mecp2^R306C/+^* (red) female mice. Mean +/- SEM; Two way ANOVA, main effect of treatment F(1, 18) = 3.916, p=0.0613; main effect of genotype F(1, 18) = 9.514, **p=0.0064; interaction effect of treatment and genotype F(1, 18) = 5.237, *p=0.0344. Tukey’s post hoc: vehicle *Mecp2^+/+^* versus *Mecp2^306C/+^*, **p=0.0066; *Mecp2^306C/+^* vehicle versus *Mecp2^306C/+^* VU0422288, *p=0.0441. (B, C) Breath rate and minute volume were not different between genotypes. N=5-6 per genotype.

## Discussion

The discovery that mutations in MeCP2 cause RTT was made over 25 years ago; however, there is currently only one FDA approved treatment for this disorder, a 3 amino acid peptide, trofinetide, derived from insulin-like growth factor (IGF) (Collins and Neul 2022; Harris 2023; Kennedy et al., 2023; Neul et al., 2023; Abbas et al., 2024; Neul et al., 2024). Trials are also now underway for *MECP2*-based gene therapy, with promising initial reports. Additional therapeutic strategies, such as RNA editing, DNA editing, X-chromosome reactivation, read-through therapies, and oligo-therapeutics are also intense foci for potential RTT treatment and hold great promise for treatment of the disorder (Collins and Neul 2022; Palmieri et al., 2023; Lopes et al., 2024; Percy et al., 2024; Vanderplow et al., 2024).

An alternate strategy for RTT therapy is to exploit targets that are disrupted upon the loss of MeCP2. While not curative, this approach may provide important adjunctive strategies for specific domains or symptoms, such as seizures or motor impairments, and may translate to other diseases that share phenotypic or mechanistic overlap with RTT. mGlu_7_ is a G-Protein Coupled Receptor that is amenable to drug development. The receptor is expressed presynaptically in the active zones of both glutamatergic and GABAergic neurons, where it functions to decrease neurotransmitter release upon stimulation (Shigemoto et al., 1996; Dalezios et al., 2002; Somogyi et al., 2003; Niswender and Conn 2010; Gee et al., 2014; Klar et al., 2015). We have demonstrated a particularly important role for mGlu_7_ in modulating GABA release in the hippocampus at Schaffer Collateral-CA1 (SC-CA1) synapses, where mGlu_7_ antagonism prevents the induction of long-term potentiation (LTP), the molecular correlate of learning and memory (Klar et al., 2015); similar effects on LTP have been shown in the amygdala (Gee et al., 2014). Using optogenetic approaches, we have demonstrated that mGlu*_7_* activation inhibits GABA release from interneurons, disinhibiting pyramidal cells in the hippocampus to induce LTP (Klar et al., 2015). We and others have also performed extensive evaluations of *Grm7* knockout (*Grm7^-/-^*) mice, and have found that they exhibit significant impairments in cognition, learning, and response to stimulants; additionally, they develop spontaneous seizures and show abnormal repetitive movements such as hindlimb clasping (Sansig et al., 2001; Bushell et al., 2002; Fisher et al., 2020). Administration of a compound that activates mGlu_7_ prevents both seizures in mice and the development of epilepsy in a kindling model (Girard et al., 2019), while mGlu_7_ NAMs and mGlu_7_ mice encoding a modified receptor C-terminus to prevent protein/protein interactions with Protein Interacting with C-Kinase 1 (PICK1) have been reported to induce seizures (Bertaso et al., 2008; Tassin et al., 2016), suggesting that increasing mGlu_7_ activity might serve as a novel therapeutic approach for epilepsy-related disorders. Additionally, recent reports have shown that people with loss-of-functions mutations in *GRM7* exhibit neurodevelopmental symptoms such as seizures, impaired myelination, severe intellectual disability, apneas, and stereotypies (Charng et al., 2016; Reuter et al., 2017; Fisher et al., 2018; Marafi et al., 2020; Fisher et al., 2021; Song et al., 2021), which are phenotypes that overlap with RTT. Combined, these studies suggest that low levels of mGlu_7_ expression are deleterious in numerous neurological domains.

We previously profiled mGlu_7_ expression in human brain samples and showed that patients with truncating mutations in RTT expressed 60-70% lower levels of mGlu_7_ in the brain (Gogliotti et al., 2017). Using a small molecule that potentiates mGlu_7_, VU0422288, we found numerous benefits in both *Mecp2^-/y^* and *Mecp2^+/-^* animals, including improvements in hippocampal LTP, cognition, social interaction, and breathing abnormalities such as apneas (Gogliotti et al., 2017). While this initial study suggested that mGlu_7_-targeted therapy could be a promising new approach, the clinical landscape of RTT is heterogeneous, and severity is dependent upon the specific MeCP2 mutation found in an individual patient as well as other factors such as X-chromosome inactivation status (Neul et al., 2008; Cuddapah et al., 2014; Gold et al., 2024; Percy et al., 2024). For example, the R133C and R306C mutations are generally milder in disease severity (Yamashita et al., 2001; Leonard et al., 2003; Neul et al., 2008; Gold et al., 2024; Percy et al., 2024). This reality suggests that trials for RTT with any drug, or, in this case, an mGlu_7_ PAM, may need to take into consideration patient mutation as a potential modifier of treatment efficacy.

Our original mGlu_7_ profiling work was limited to seven patients, and all had truncating mutations that are associated with severe disease (Gogliotti et al., 2017). For the current study, we obtained additional samples from patients with a range of RTT mutations to profile the expression of *GRM7* (**Table 1**, (Smith et al., 2022)). Additionally, although we previously showed that the activity of the group III mGlu receptor PAM VU0422288 was mediated by mGlu_7_ using pharmacological tools to exclude mGlu_4_ and mGlu_8_, we also profiled the expression of *GRM4* and *GRM8* across this wider cohort. These studies revealed that, among the three widely expressed group III mGlu receptors, only *GRM7* mRNA was significantly reduced. We would note that there were three samples that were ranked as outliers via a ROUT test (**Table 1, starred**), and we currently do not know why these samples are outliers as they span several different mutations (R255X, M246Del, and mutation-negative, **Table 1**). We did not observe a correlation in expression between PMI and expression, and, although we did observe significant negative correlations of *GRM7* and *GRM8* expression with age in the control group, this was not the case for RTT. Currently, we do not have an explanation for these differences in *GRM7* mRNA expression for these specific RTT patients, although X-chromosome inactivation status or medication usage may contribute. We are limited in this interpretation as medical records were not available for these de-identified patient samples.

Interestingly, while we did not establish any correlation to suggest co-regulation of *GRM7* with *GRM4* or *GRM8*, we found a very strong correlation of the expression of *GRM4* and *GRM8* in both control and RTT groups (**Figure 3**). This finding is intriguing from the standpoint of mGlu receptor heterodimerization. While the mGlu receptors function as constitutive dimers, they have recently been found to heterodimerize (Doumazane et al., 2011; Yin et al., 2014; Levitz et al., 2016; Moreno Delgado et al., 2017; Habrian et al., 2019; Lee et al., 2020; Xiang et al., 2021; Lin et al., 2022; Kukaj et al., 2023; Lin et al., 2024). mGlu_4, 7 and 8_ all belong to the group III mGlu receptor subfamily, and can heterodimerize among each other as well as with the group II mGlu receptors, mGlu_2_ and mGlu_3_ (Doumazane et al., 2011; Lee et al., 2020). mGlu receptor heterodimers represent a fascinating pharmacological challenge and opportunity, as drug development for the receptors has, thus far, proceeded with homodimer-expressing systems and emerging evidence indicates that heterodimers exhibit dramatically different pharmacology compared to homodimers (Yin et al., 2014; Liu et al., 2017; Moreno Delgado et al., 2017; Lee et al., 2020; Xiang et al., 2021; Lin et al., 2022; Lin et al., 2024). The observation that *GRM4* and *GRM8* may be co-regulated at the mRNA level in the human brain suggests that this receptor combination may be particularly important from a heterodimer standpoint. Additional co-localization studies will surely shed light on this interesting and robust correlation, and single cell studies will be needed to understand if this correlation reflects co-expression in individual cells, such as neurons, or among cells of different types.

When RTT samples were binned into “severe” and “mild” mutations, we found that the cohort of patients with the R168X, R255X and R270X mutations exhibited significant reductions in *GRM7* mRNA and protein compared to patients with historically mild mutations. Interestingly, we also quantified a significant decrease in expression in patients who had been clinically diagnosed with RTT, but for whom we were unable to detect a mutation in *MECP2* by Sanger sequencing. There are several other disorders that exhibit strong phenotypic overlap with RTT, and patients are often diagnosed with RTT until the causative gene is identified. Some of these other “RTT-like” genes including Transcription Factor 4 (*TCF4*, the causative gene for Pitt Hopkins Syndrome ) (Wirgenes et al., 2012; Sweatt 2013; Kennedy et al., 2016), Cyclin-Dependent protein Kinase Like 5 (*CDKL5*), which causes *CDKL5* Deficiency disorder (Szafranski et al., 2015; Leonard et al., 2022), and Forkhead box G1 (*FOXG1*), which causes *FOXG1* syndrome (Brockmann and Staudt 1993; Mencarelli et al., 2010; Akol et al., 2022; D’Mello 2023). While we are unsure if the patient samples used here exhibit mutations in these genes, we are currently performing studies in a *Tcf4^+/-^* model examining mGlu_7_ expression and the therapeutic potential of PAMs which may shed light on the utility of mGlu_7_ potentiation in other disorders related to RTT. Additionally, while we have not determined the mechanism of mGlu_7_ dysregulation in RTT, we hypothesize that one mechanism may be via microRNAs (miRNAs). The Gogliotti lab has recently performed a profiling study of miRNAs in a subset of the samples using for profiling here (Vanderplow et al., 2024). These studies have identified numerous microRNAs that are dysregulated in RTT autopsy samples; one of these is miR-15a, which has been shown to regulate mGlu_7_ expression in the context of schizophrenia (Beveridge et al., 2010). Future studies will determine if mGlu_7_ expression is regulated by this *MECP2*-impacted miRNA and how this regulation extends, or does not, to other disorders with RTT overlap.

Our correlations of mGlu_7_ expression with mutations that induce higher levels of clinical severity suggested that it was possible that our previous findings with VU0422288 in *Mecp2^+/-^* mice could be limited to the context of specific *MECP2* mutations. To test this hypothesis, we profiled mGlu_7_ protein expression in R306C mice. These studies showed that mGlu_7_ was not reduced in synaptosomes prepared from the cortex, cerebellum, or brainstem of these animals compared to controls. In contrast, mGlu_7_ levels were significantly reduced in the hippocampus from *Mecp2^R306C/+^* mice. This may suggest that the effects of *Mecp2* loss or mutation are brain region or cell-type specific as it relates to mGlu_7_ expression, and our human studies in the cortex may not reflect changes in other brain areas such as the hippocampus. The lack of decrease in the brainstem and cortex suggested that it was possible that *Mecp2^R306C/+^* mice might not respond to VU0422288 in terms of apneas, a robust phenotype we had observed was responsive to VU0422288 in *Mecp2^+/-^* animals (Gogliotti et al., 2018). Encouragingly, we did find a significant reduction in apneas when *Mecp2^R306C/+^* animals were dosed with VU0422288. These results suggest, again, that there may be cell-type specific effects on expression that were not captured in our synaptosomal preparations; however, they also suggest that mGlu_7_ PAMs may have broad utility in patients with a variety of RTT mutations, at least for specific phenotypes. Systematic studies are needed across models of multiple MeCP2 mutations to truly address this possibility. Additionally, there may be utility in the use of an mGlu_7_-specific imaging agent to understand mGlu_7_ expression across the entire brain in RTT to validate the therapeutic potential of mGlu_7_ PAMs in multiple aspects of the disease phenotype.

## Authorship contribution statement

**Geanne A. Freitas:** Writing – review & editing, Visualization, Methodology, Investigation. **Sheryl Anne D. Vermudez:** Writing – original draft, Visualization, Methodology, Investigation. **Mackenzie Smith:** Writing – review & editing, Visualization, Methodology, Investigation. **Rocco Gogliotti:** Writing – review and editing, Visualization, Methodology, Investigation, Supervision, Resources, Project administration, Methodology, Funding acquisition, Formal analysis, Conceptualization. **Colleen M. Niswender:** Writing – review & editing, Writing – original draft, Supervision, Resources, Project administration, Methodology, Funding acquisition, Formal analysis, Conceptualization.

## Data availability

All mentioned data are presented in this published article or the supplementary information or are available from the corresponding author upon reasonable request.

## Ethics statement

This manuscript is an original work and is not under consideration for publication elsewhere. All authors have reviewed and approved of the submission.

## Funding

This work was supported by grants R01MH124671, R01NS031373, R21MH102548, R01MH113543, K01MH112983, R01NS112171, and the International Rett Syndrome Foundation. Harvard Brain Tissue Resource Center is supported by Public Health Service contract HHSN-271-2013-00030, and the University of Maryland Brain Bank is a tissue repository of the National Institutes of Health (NIH) NeuroBioBank.

## Declaration of Competing Interest

The authors declare that the research was conducted in the absence of any commercial or financial relationships that could be construed as a potential conflict of interest.

## Acknowledgements

We thank the William K. Warren Foundation for the endowment of the Warren Center for Neuroscience Drug Discovery.

## Notes

### Competing Interest Statement

The authors have declared no competing interest.

