## Supplemental Figures 1-4 for "Profiling metabotropic glutamate receptor 7 expression in Rett syndrome: consequences for pharmacotherapy"

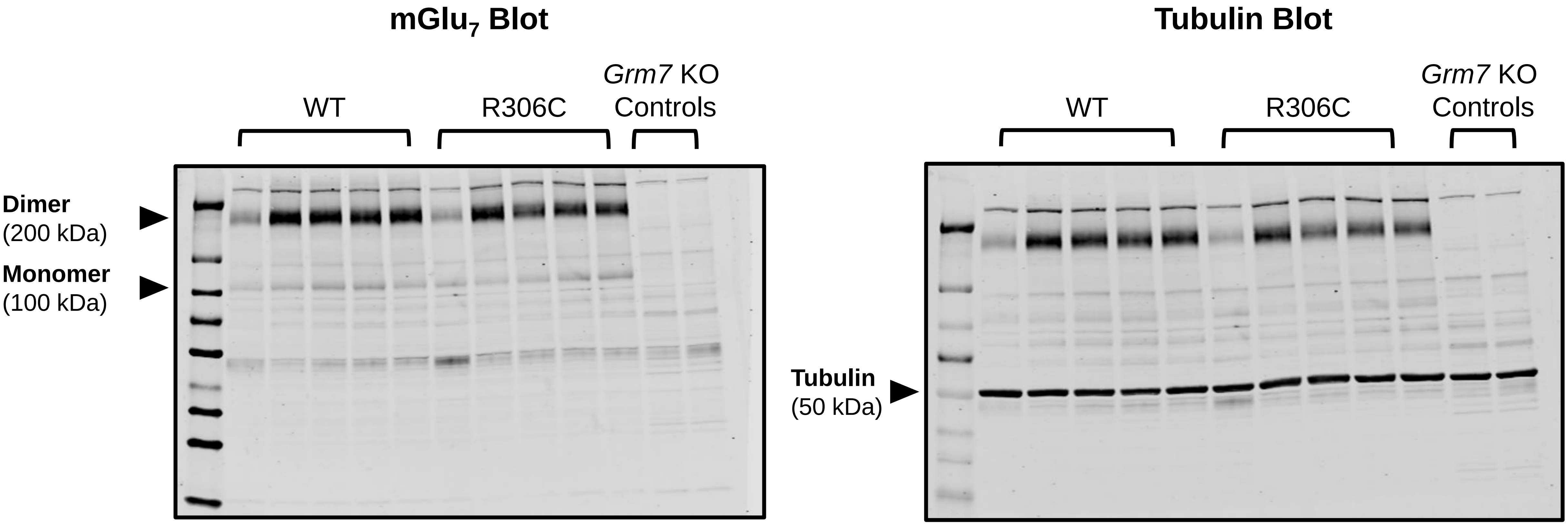


**Supplementary Figure 1. Complete Western blots of *Mecp2*$^{R306C/+}$ mice hippocampal synaptosome samples.**

Immunoblots of mGlu$_7$ in the hippocampus of *Mecp2*$^{R306C/+}$ mice synaptosomes. Bands corresponding to the mGlu$_7$ dimer and monomer are shown. Tubulin (50 kDa) was used as loading control. n = 5 mice per genotype.

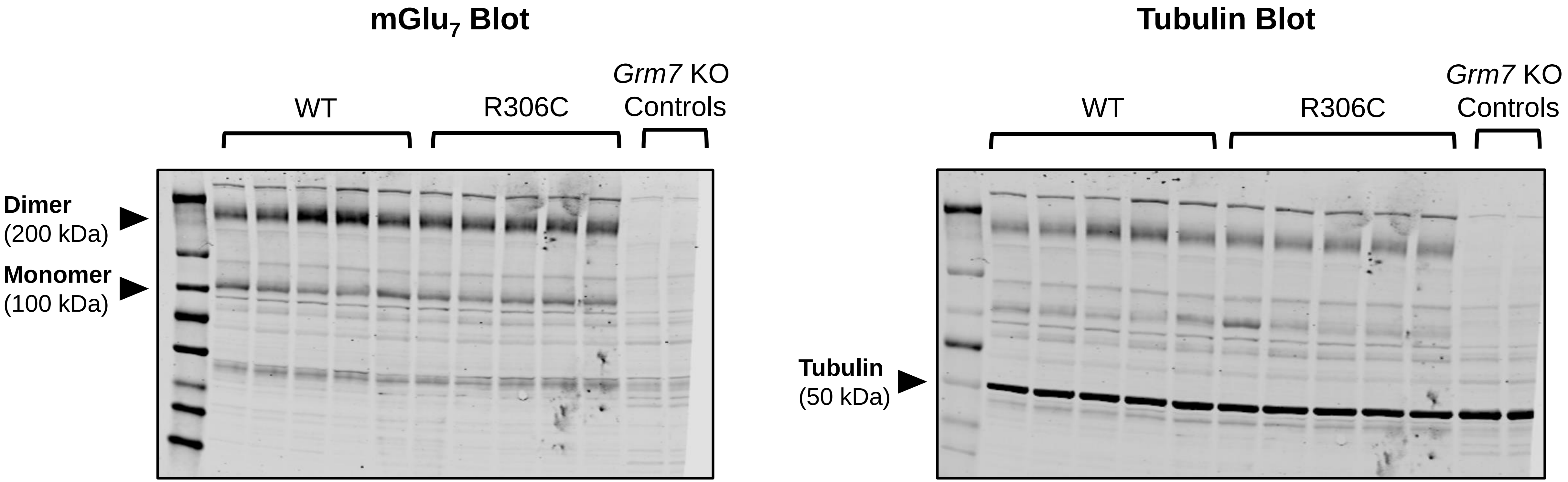


**Supplementary Figure 2. Complete Western blots of *Mecp2*$^{R306C/+}$ mice cortical synaptosome samples.**

Immunoblots of mGlu$_7$ in the Cortex of *Mecp2*$^{R306C/+}$ mice synaptosomes. Bands corresponding to the mGlu$_7$ dimer and monomer are shown. Tubulin (50 kDa) was used as loading control. n = 5 mice per genotype.

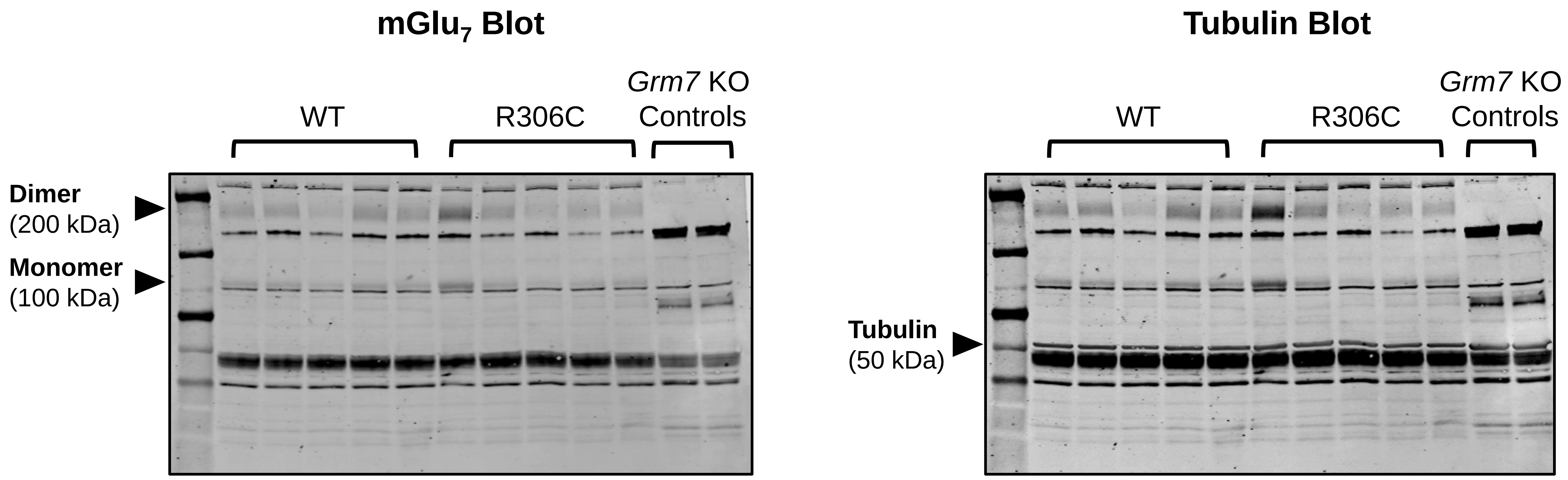


**Supplementary Fig. 3. Complete Western blots of *Mecp2*$^{R306C/+}$ mice cerebellar synaptosome samples.**

Immunoblots of $mGlu_7$ in the cerebellum of *Mecp2*$^{R306C/+}$ mice synaptosomes. Bands corresponding to the $mGlu_7$ dimer and monomer are shown. Tubulin (50 kDa) was used as loading control. n = 5 mice per genotype.

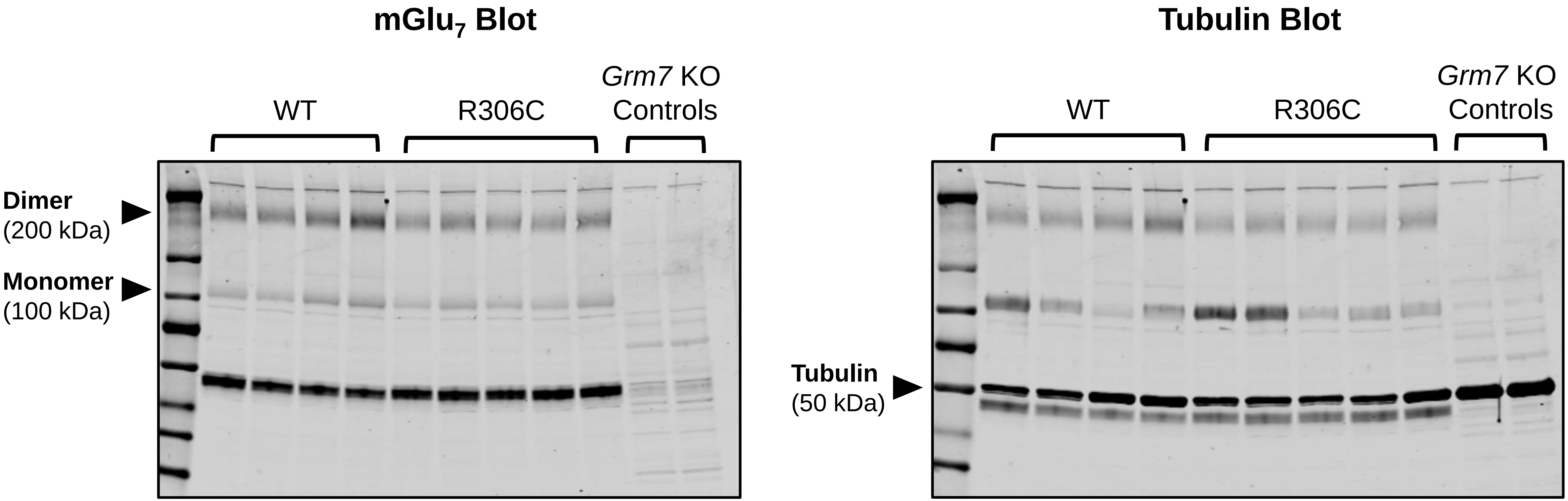


**Supplementary Fig. 4. Complete Western blots of *Mecp2*$^{R306C/+}$ mice Brainstem synaptosome samples.**

Immunoblots of $mGlu_7$ in the brainstem of *Mecp2*$^{R306C/+}$ mice synaptosomes. Bands corresponding to the $mGlu_7$ dimer and monomer are shown. Tubulin (50 kDa) was used as loading control. n = 5 mice per genotype.
